# Noise statistics drive background- and channel-specific interference in auditory midbrain representations of speech

**DOI:** 10.64898/2026.06.30.735623

**Authors:** Jimmy Dion, Jenna Blain, Ruotong Li, Ian H. Stevenson, Monty A. Escabí

## Abstract

Although humans and animals excel at interpreting target sounds in competing noise, sound recognition abilities can vary widely due to statistical characteristics of the interfering background. Yet, how the brain leverages statistical sound information to segregate sounds in noise remains unclear. Here, using many natural auditory textures as background stimuli, we test how the encoding of speech is altered by natural noises in the inferior colliculus (IC) of unanesthetized rabbits. We identify foreground- and background-driven neural response components to sound mixtures and find that the background statistics alter the neural representation of speech in a frequency- and modulation-specific manner. Neural encoding signal-to-noise ratios (SNRs) vary extensively across background categories, frequencies, and modulations, and these influence the encoding of speech independently of the acoustic SNR. This variability is driven by both the background spectrum and modulation statistics, which distort the speech representation and show distinct interference effects in the modulation ranges for rhythm and pitch. Thus, spectrum and modulation statistics critical to speech recognition in noise are reflected in IC neural population activity. These neural response statistics and resulting distortions likely determine the coding fidelity for speech in downstream cortical regions and provide a neural basis for differences in speech intelligibility in real-world noises.

## Introduction

In most listening scenarios, background environmental noises compete with sounds of interest for a listener’s attention, and the acoustic properties of the competing noise mask or distort the target sounds. Yet, humans and animals can successfully recognize target foregrounds, such as speech or vocalizations, and segregate them from noisy surroundings ^1^. For typically-hearing people, speech is often intelligible even when the power of the noise is considerably greater than the power of the speech with signal-to-noise ratios (SNRs) as low as -10 dB. How the auditory system achieves this task, and how listening in noise can be improved for people with hearing loss are ongoing research questions ^2,3^. As sound pressure waveforms add linearly and are transduced into neural activity patterns by the cochlea, mixtures of speech and noise are intertwined into a single peripheral neural representation. How the downstream auditory regions segregate speech when embedded in noise under different and changing backgrounds is not well understood. Here, using neural responses in the rabbit inferior colliculus to speech mixed with natural and perturbed noises, we characterize how speech representations vary between backgrounds and at multiple acoustic SNRs.

The statistics of background noise play a critical role in determining whether and when speech perception is impaired. *Energetic masking* or spectrum-based interference occurs when the spectral power distribution of a background sound overlaps with that of the target foreground sound ^3–6^. Beyond energetic masking, modulation masking, driven by spectro-temporal modulation statistics of natural sounds, creates a secondary source of interference that can either help ^7–10^ or hinder ^11–14^ the recognition of a foreground signal. For speech recognition, these two forms of acoustic interference lead to highly variable masking outcomes where, depending on the background, recognition accuracy can vary from near perfect to near chance, even when the mixtures are presented at the same overall acoustic SNR ^10^. Since differences in recognition accuracy can be directly attributed to the spectrum and modulation statistics of each individual noise, neural representations of speech may show similar channel-specific interference.

Previous work suggests that the brain performs a multi-level, hierarchical decomposition of natural sounds that is essential for sound recognition and for separating speech from noise^15,16^. After the cochlea decomposes sounds into frequency components, brainstem and midbrain circuits appear to decompose sounds into detailed spectro-temporal modulation components ^17–19^. Thalamic and cortical circuits further decompose sounds into coarser resolution modulations ^20,21^ and sparse spectro-temporal features that are noise-invariant ^16,22^. Despite substantial changes in spectro-temporal selectivity along the auditory pathway, the central nucleus of inferior colliculus (ICC) is the first nucleus where modulation tuning predominates ^23^. Distortion of the neural representation of speech in the ICC is thus likely to be a key determinant of speech-in-noise perception.

Here, using a diverse set of natural background sounds and their perturbed variants, we characterize the neural representation of speech-in-noise in the rabbit ICC. We develop a shuffling approach to separate the foreground-driven and background-driven power in individual neural frequency and temporal modulation bands and then measure the midbrain SNR (mSNR) across these dimensions. We find that speech and natural background sounds are jointly-encoded at the level of the ICC. Both the spectrum and modulation statistics of natural background sounds distort the neural representation of speech by exerting opposing influences that improve or distort the representation of word- or articulatory-level speech features. However, we find that the mSNR patterns occur mostly independently of the overall acoustic SNR, suggesting that the unique statistics of each individual background distort the speech representation consistently regardless of the overall noise levels. Altogether, these findings suggest that the spectrum and modulation statistics of different background sounds lead to feature-specific alterations in the coding of speech within the inferior colliculus. Considering these feature-specific neural SNR patterns may be necessary to explain differences in speech-in-noise perception beyond the effect of the overall acoustic SNR.

## Results

We recorded neural population activity in the ICC of head-restrained unanesthetized Dutch belted rabbits (n=5) presented with speech in competing sources of environmental noise. The ICC receives convergent input from brainstem structures, and neurons within the ICC are selective for spectro-temporal modulations that are critical features in natural sounds ^18,19,24–26^. Both the spectrum and modulation statistics of natural background sounds strongly influence human perception of speech^10,27^, and here we determine how such natural sound statistics interfere with the neural representations of speech in the ICC.

### Neural Response Statistics in the ICC Reflect the Statistical Structure of Speech

Animals passively listened to either clean speech (isolated foreground with no background) or speech (four excerpts from an audiobook version of Shakespeare’s *Hamlet*, read by an adult male speaker) mixed with original and perturbed variants of eleven background sounds. Collectively, these sounds cover a range of spectrum and modulation statistics present in real-world listening environments that strongly impact human speech recognition performance in noise ^10,27^. Below, we first focus on characterizing neural responses to clean speech, since these provide a reference to study responses to speech mixed with different natural background sounds.

Here we analyze neural signals on each recording channel using the analog multi-unit activity (aMUA, see Methods). Due to the tonotopic organization of the ICC ^28^, frequency response areas for the aMUA signal on each channel along the linear probe (Fig. 1A) exhibit a clear gradient. For the typical recording site shown here, the gradient extends between ∼0.5 to 9 kHz. In response to clean speech, neural population activity exhibits a synchronized and highly-structured spatio-temporal pattern (Fig. 1B). Coarse, speech-driven temporal fluctuations (<10Hz) are evident that follow the amplitude modulations in speech within and between consecutive words. Also evident are faster temporal fluctuations (∼90-500 Hz) that are synchronized to fine temporal details in the speech signal. Differences in the activity across recording channels likely reflect differences in overlap between the spectral tuning of each channel and the power spectrum of speech, with the most-active recording channels overlapping sound frequencies with high relative power.

**Figure 1:**
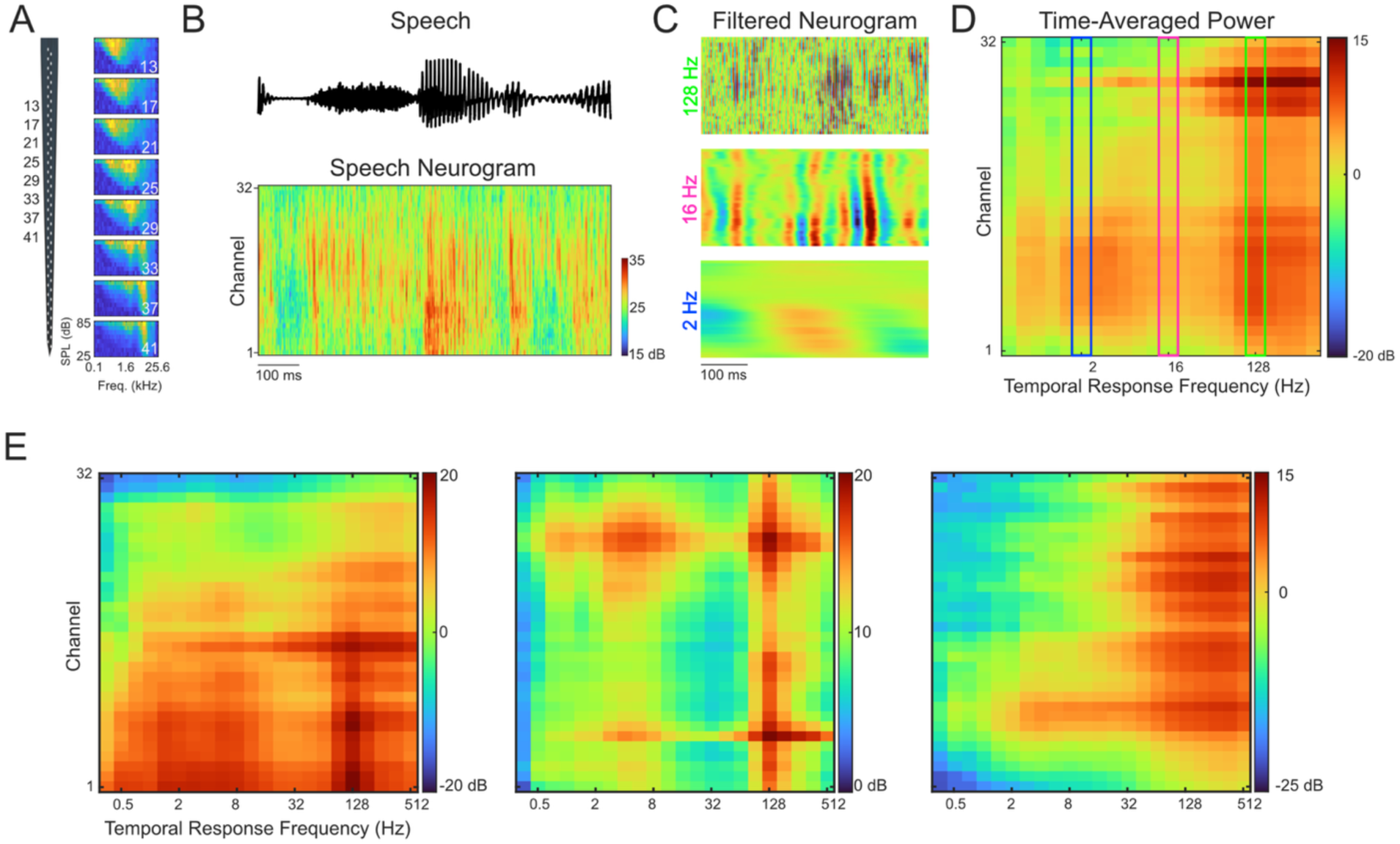
Typical multi-unit ICC responses to clean speech. (A) Frequency response areas for one example recording site for evenly-spaced channels along the linear probe. Responses to tone pips varying in frequency and level illustrate the tonotopic organization of ICC. (B) Analog multi-unit activity (aMUA) across channels in the recording array (the neurogram, bottom) in response to one exemplar of clean speech (top) for the same example recording site. (C) Log-spaced octave-wide temporal bandpass filters act on each channel, decomposing the neurogram into temporal modulation subbands. Here, outputs for three filters are shown with center frequencies of 2, 16, and 128Hz, respectively. (D) The midbrain modulation power spectrum (mMPS) calculated by time-averaging the cross-trial power for each channel and temporal modulation subband, is shown here averaged over all pairs of clean speech excerpts and trials. (E) mMPS for three additional example recording sites. dB scales indicate power relative to 1uV.

To determine to what extent modulations in the neural population response patterns reflect the modulation structure in speech, we first estimate a midbrain modulation power spectrum (mMPS). Specifically, we decompose the aMUA signals on each channel - the neurogram - using a bank of temporal filters spanning 0.25 to 512 Hz (Fig. 1C). Since aMUA signals are generated by filtering and envelope extraction, the response of each temporal filter indicates envelope fluctuations within the neural signal. Additionally, since aMUA signals reflect a filtered, band-limited response to the original sound, we can compare the output of these temporal filters directly to temporal modulations present in the sound. Here, the neurogram outputs for the low-frequency modulation subbands represent coarse fluctuations in the neural response, while higher subbands capture neural responses with successively more temporal detail. Following this multi-resolution decomposition of the neurogram, the mMPS is then computed by measuring the cross-trial synchronized power for each electrode channel and each modulation subband (Fig. 1D, see Methods). Similar to previous methods that use shuffling to estimate stimulus- and noise-driven correlations, this approach removes the contributions of noise to the modulation power (see Methods). The mMPS is motivated by the modulation power spectrum statistic of natural sounds, which represents the acoustic power as a function of the sound frequency and temporal modulation frequency^29^. However, where the conventional MPS uses an auditory model decomposition to characterize the statistics of natural sounds, the mMPS is derived directly from neural activity and reflects the statistics of ICC neural responses.

The resulting mMPS measures the power of synchronized stimulus-evoked neural activity as a function of both the electrode channel and the temporal response frequency. For this example, recording site (Fig. 1D), there is a strong mode of stimulus-evoked activity for temporal frequencies above ∼128 Hz, with only a subset of low-frequency channels showing stimulus-evoked activity for lower temporal modulations <16Hz.

Synchronized activity to speech varies across recording locations. However, across recording sites, we find that the mMPS tends to have modes that are generally localized to the rhythm (<16 Hz temporal modulation frequency) and the pitch or voice periodicity ranges (>100 Hz). Midbrain modulation power in both of these bands likely indicates neural populations that contain information reflecting both relatively slowly-varying speech features such as transitions between words and phonemes, as well as articulatory features such as pitch and formants. Speech has prominent fluctuations below 10 Hz, at the timescales of words and phonemes, that are critical for word identification and recognition ^30,31^. Speech also has substantially faster periodicities (> 100 Hz) that are generated by vocal fold vibration and that likely contribute to pitch perception, voice identification, and source segregation ^30,32,33^. In ICC we find that certain recording sites exhibit a stronger mode of synchronized activity exclusively within the voice periodicity range (Fig. 1E, right), while other sites display bimodal synchronized responses that also occur at slower rhythmic fluctuations (Figs. 1, left and middle). After aligning recording sites based on the characteristic frequency (CF) of each channel (see Methods), the bimodal neural population response is also evident in the clean speech mMPS averaged across recording sites (Fig. 2A).

**Figure 2:**
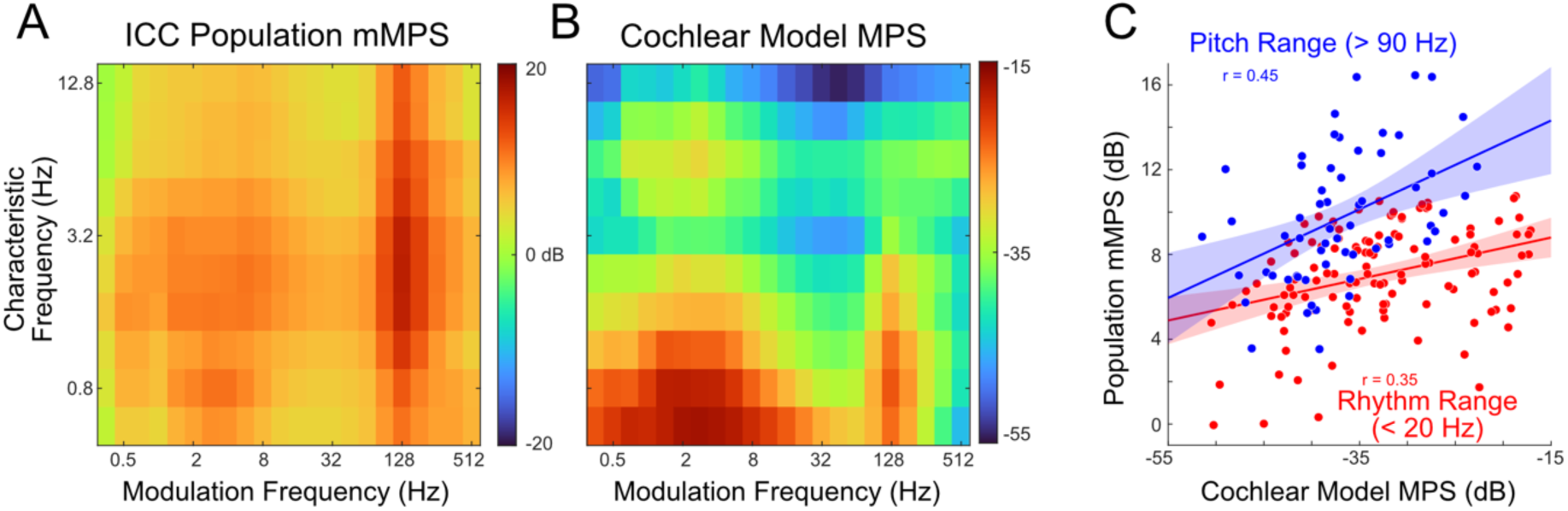
Population-average ICC midbrain modulation power spectrum (mMPS) for clean speech and comparison with a cochlear model. (A) Average mMPS across recording sites (n=41), generated by aligning recording channels by their characteristic frequency. dB indicates power relative to 1uV. Results indicate cross-trial power and are averaged over excerpts and trials. (B) Modulation power spectrum for a cochlear model. dB indicates power relative to 1 (standardized waveform units). (C) Modulation power in ICC mMPS is correlated with the cochlear model power, here analyzed separately for the pitch range (>90 Hz modulation frequency) and rhythm range (<20 Hz modulation frequency). Dots denote pairings of frequency (range: 0.6 – 18 kHz) and temporal modulation (rhythm range in red and pitch range in blue). Lines denote linear fits on the dB scale. Bands denote 95% confidence intervals for the fits.

To determine the extent to which the mMPS directly reflects the modulation structure in speech, we used a peripheral auditory model decomposition to estimate the modulation power spectrum (MPS) of the speech sounds (see Methods). As with the experimentally-observed ICC activity, the speech model MPS is dominated by two distinct modes in the ranges of ∼2 Hz and ∼128 Hz temporal modulation frequency (Fig. 2B). The latter is consistent with the talker’s voice periodicity, since clear periodic voice fluctuations occur in the sound waveform at the same frequencies (100-170 Hz) ^25,30,34^. However, unlike IC neural activity (Fig. 2A), which synchronized broadly to these components irrespective of the recording site characteristic frequencies the synchronized modes in the model MPS occur more strongly at low-frequency channels (<1kHz). Nonetheless, the observed response patterns for the slow (<20 Hz) and fast (>90 Hz) subbands were significantly correlated between the ICC data and the model MPS (r=0.35; p<0.001 for rhythm range, r=0.45; p<0.001 for pitch range).

Collectively, these results illustrate that sound-synchronized spatio-temporal activity patterns in the ICC reflect statistics of spectro-temporal features in speech, including both slow fluctuations that are generated by transitions of words and phonemes^30,31,35^, as well as finer-detailed spectro-temporal features such as voicing periodicity and fine structure ^25,30,34^.

### Measuring Background Noise Interference with Speech Representations

The results above demonstrate that ICC neural activity reflects statistical features of the acoustic signal at multiple spectro-temporal resolutions in response to clean speech. Here we examine how different background noises, each with unique spectrum and modulation statistics, distort the neural representation of speech. We presented mixtures of speech combined with 11 background noises, including eight-speaker babble, birds, fire, water, and white noise, at a constant acoustic signal-to-noise ratio (aSNR) of +3 dB. Even at a constant aSNR, these background sounds have previously been shown to produce highly-variable speech recognition accuracy for human listeners ^10,27^. Here, we use the cochlear model output to illustrate the different statistics of the foreground speech and the backgrounds (Fig. 3, shown for eight-speaker babble). Both the speech and backgrounds contain rich spectro-temporal structure, and in the case of some backgrounds, such as eight-speaker babble, the foreground and background statistics are similar. Both the clean speech and eight-speaker babble exhibit relatively slow and coherent time-frequency fluctuations in power dominated by frequencies below 1kHz (Fig 3, bottom row for clean speech and original babble-8). The dominant spectro-temporal modulations for both sounds are relatively slow (<10 Hz temporal modulation frequency) and have relatively broad spectral features (<3 cycles/octave spectral modulation frequency).

**Figure 3:**
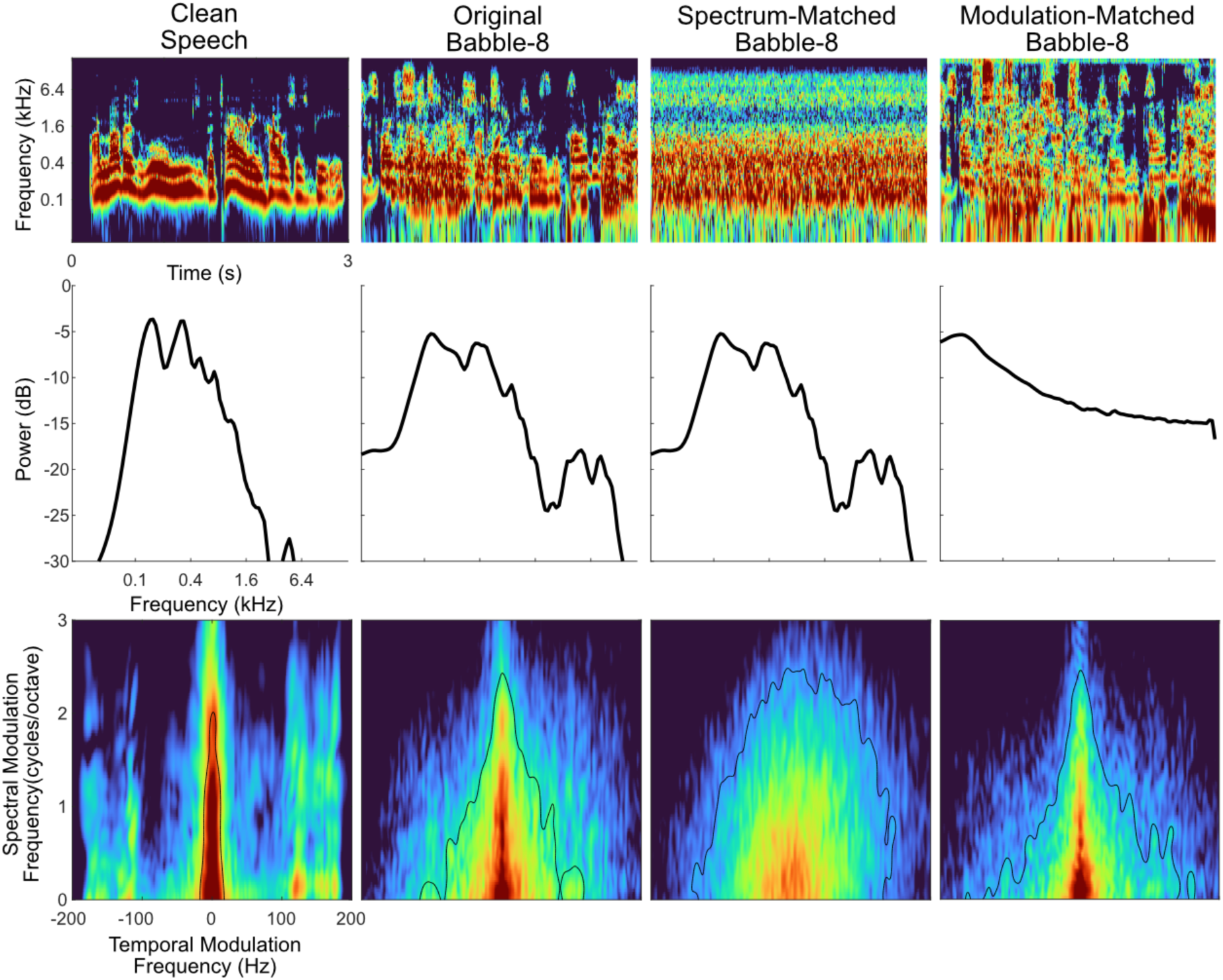
Sound statistics for speech and noise. Cochlear model output (top) for speech (left) and eight-speaker babble backgrounds (right), along with power spectra (middle) and modulation power spectra (MPS, bottom) for each sound. Synthetic backgrounds that are spectrum-matched (SM) or modulation-matched (MM) were generated by phase randomizing in the Fourier domain (SM) or equalizing the spectrum to 1/f (MM). This allows us to isolate the contribution of modulation and spectrum statistics, respectively. Black lines in MPS denotes 90% power contours.

To determine how the spectrum and modulation statistics of these background sounds impact neural encoding in the ICC, we use two manipulations to generate synthetically-perturbed backgrounds (see Methods). First, we whiten the modulation statistics while preserving the power spectrum leading to a spectrum-matched (SM) background (Fig. 3). Second, we manipulate the spectrum statistics by equalizing the power spectrum to follow a 1/*f* trend. This produces approximately constant power for cochlear filters, whitening the sound spectrum while preserving the MPS, thus producing a modulation-matched (MM) background sound (Fig. 3). Together, the SM and MM backgrounds isolate the influence of spectrum statistics from that of modulation statistics on speech-driven neural activity in the ICC.

ICC population responses to speech-in-noise mixtures synchronize to the combined acoustic features in the mixture and reflect the interference between foreground and background sounds. To identify the foreground- and background-driven response components, we developed a procedure based on a block design where speech and background sound mixtures are generated from a set of 4 foreground excerpts and 4 background excerpts (Fig. 4A). Combining these excerpts produces 16 mixtures that are then delivered pseudo-randomly, and responses are then analyzed by shuffling across excerpts with either the foreground speech or the background kept frozen (see Methods).

**Figure 4:**
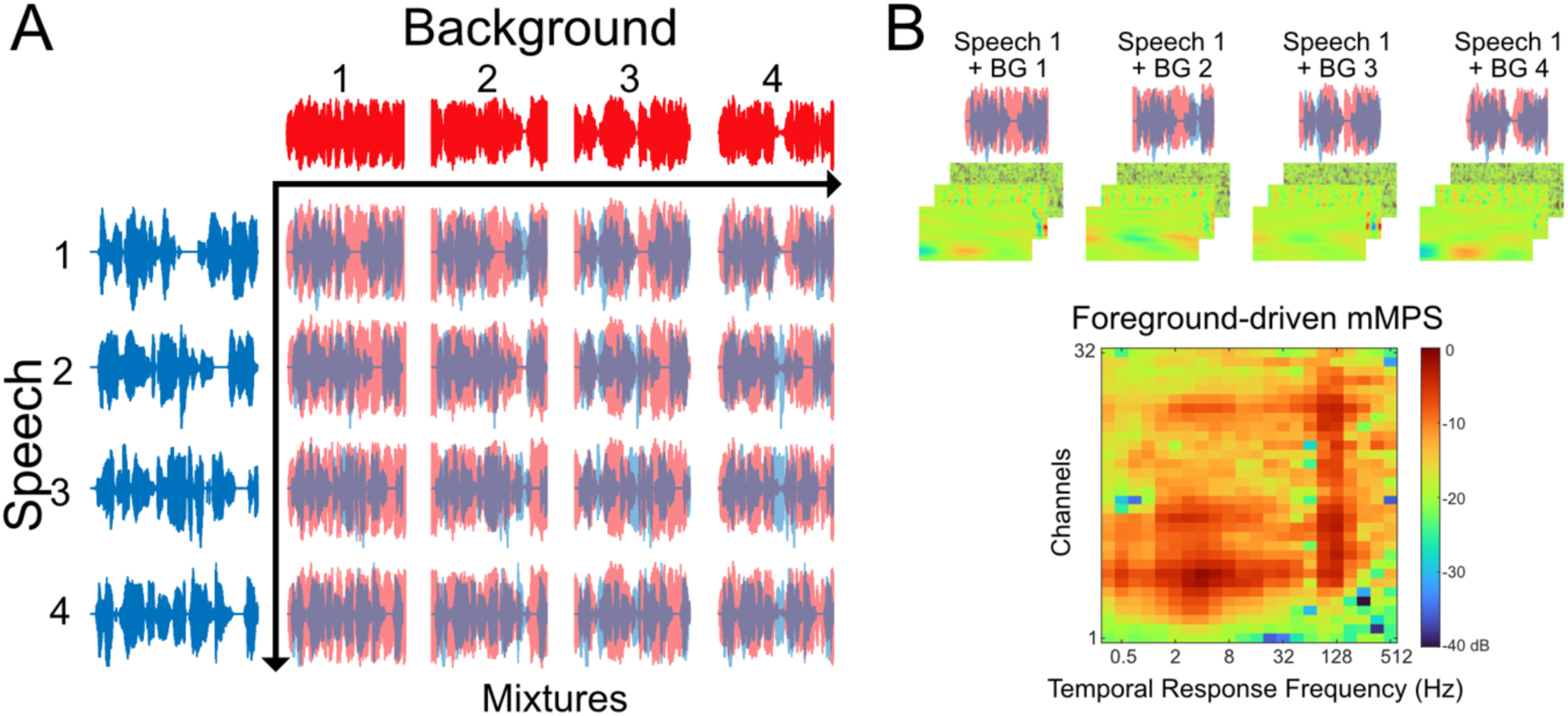
Estimating foreground- and background-driven modulation power spectra by shuffling across background and foreground excerpts. A) Speech-in-noise mixtures are generated by combining one of four speech excerpts (blue) with one of four background excerpts (red), leading to 16 unique mixture stimuli. (B) To estimate the foreground-driven mMPS, response neurograms for each of the sound mixtures are filtered and decomposed into subbands (B; similar to Fig. 1C). The cross-trial power is then computed for a frozen speech excerpt (different background excerpts), followed by averaging over background excerpts. The resulting foreground-driven mMPS is shown here for one example recording site, with original bird chorus background excerpts. To estimate the background-driven mMPS (not shown), we follow the same procedure but calculate the cross-trial power with a frozen background excerpt and variable speech excerpts.

Building on the cross-trial mMPS calculation above and previous work using shuffling to isolate stimulus and noise correlations ^24,36^, our approach here is to shuffle across subsets of mixtures to isolate the foreground or background driven mMPS. By computing the subband-filtered synchronized power *between* multiple background excerpts for *frozen* foreground excerpt (Fig. 4B) and averaging across foregrounds (see Methods), we can estimate the foreground-driven mMPS (mMPS_F_). Since the foregrounds are fixed and the backgrounds differ across shuffled trials, the trial-shuffled mMPS estimate contains only the common foreground-driven mMPS (Fig. 4B, bottom). Similarly, the background-driven mMPS (mMPS_B_) is estimated by computing the subband-filtered synchronized power *between* foreground excerpts for a *fixed* background excerpt.

Typical foreground- and background-driven mMPS estimates are quite distinct from each other, despite being calculated from the same set of mixture stimuli (Fig 5). The foreground speech-driven mMPS generally mirrors the bimodal patterns seen in the mMPS for clean-speech (Fig. 1), with distinct peaks at both fast modulations near ∼128Hz temporal modulation frequency as well as slow modulations <20Hz. The background-driven mMPS, by contrast, is more broadly distributed across many electrode channels, with synchronized activity between 1 - 256 Hz.

**Figure 5:**
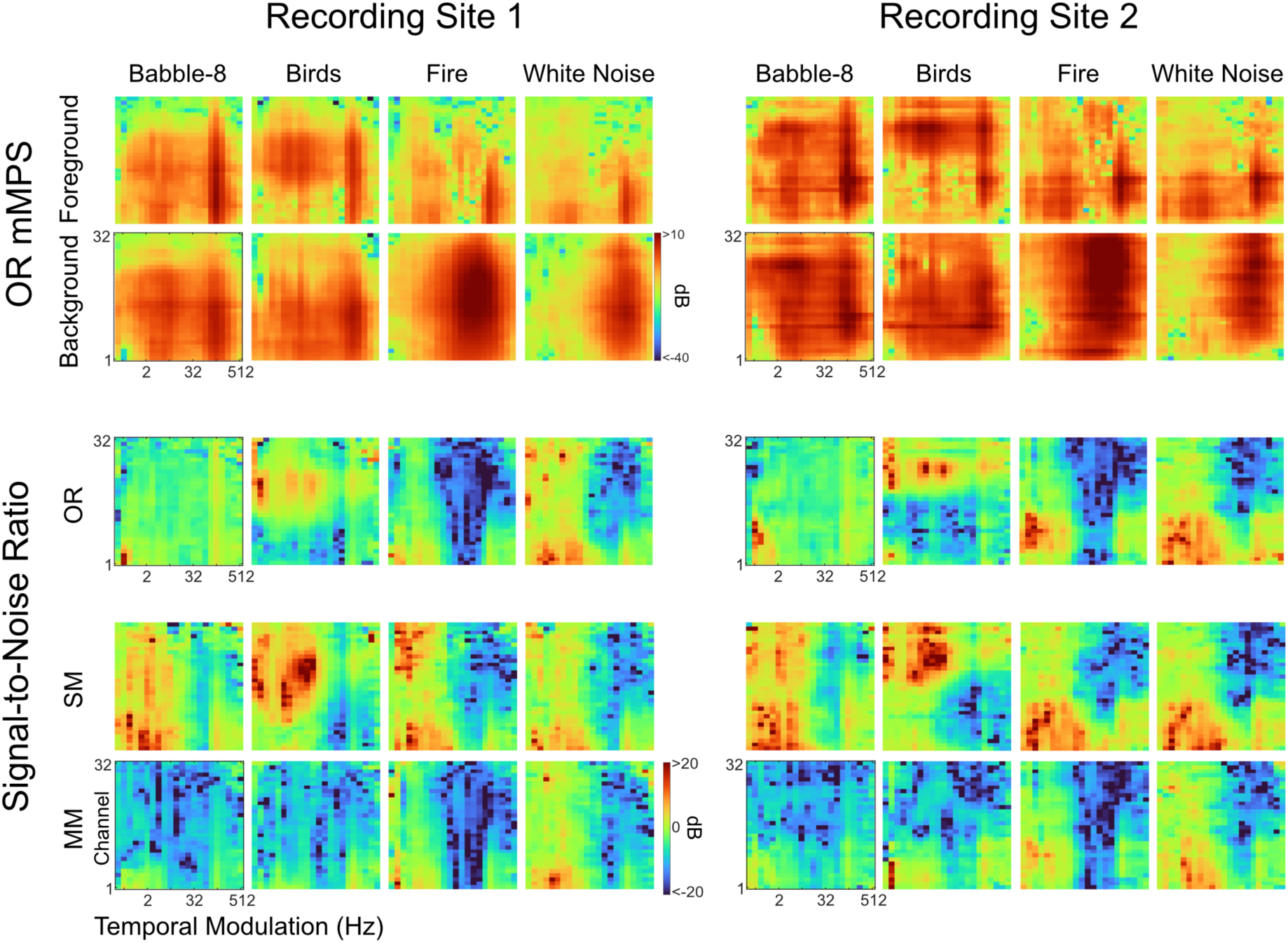
Foreground- and background-driven mMPS (top two rows) and neural signal-to-noise ratios (bottom three rows). Data from two example recording sites with the background- and foreground-driven mMPS for sound mixtures of speech and eight-speaker babble, bird chorus, fire, or white noise (top). The ratio of these two power estimates gives a neural signal-to-noise estimate for each channel and modulation subband. Here, we show SNR for speech mixed with original backgrounds, as well as the spectrum-matched (SM) and modulation-matched (MM) synthetic backgrounds (bottom).

We find that the background-driven mMPS for each recording site varies considerably across backgrounds (Fig. 5, second row). Synchronized responses occur across different subsets of recording channels and cover different temporal modulation ranges. For instance, eight-speaker babble produces a strong bimodal pattern of synchronized neural activity in the rhythm and voice periodicity ranges, similar to that observed for clean speech. In contrast, responses to white noise and fire lack synchronization in the rhythm range and instead appear to synchronize strongly only to temporal modulation frequencies >16 Hz. These differences in synchronized activity across background sounds are consistent with the fact that temporal modulations present in these backgrounds vary considerably and that IC neurons are known to faithfully synchronize to spectro-temporal sound modulations^18,25,37^.

Additionally, the foreground-driven mMPS appears to be modulated by the interfering background sounds (Fig. 5, top row). While the foreground speech-driven activity in mixtures of speech and eight-speaker babble closely resembles the pattern observed for clean speech, speech-driven synchronized activity shifted towards deeper recording locations (higher characteristic frequencies) for speech mixed with bird babble. Conversely, speech-driven synchronization shifted towards shallower recording locations (lower characteristic frequencies) when speech was mixed with white noise. Comparing across recording sites, we find that the mMPS_F_ and mMPS_B_ are often qualitatively similar but shift up or down depending on the frequency range covered by the recording site. Although the relative channel numbers change, mMPS estimates show consistent trends across recordings (Fig 5, left vs right), and these results demonstrate that the foreground speech and background sounds are represented jointly in IC responses to mixtures. The differences in the mMPS_F_ across backgrounds suggest that the background- and foreground-driven neural activity are not strictly additive.

To quantify how individual background sounds impact the fidelity of speech encoding, we use the mMPS_B_ and mMPS_F_ for each recording site to estimate a midbrain signal-to-noise ratio (mSNR; see Methods). The mSNR quantifies how speech-synchronized neural activity is affected by the interfering background as a function of both the electrode channel location (or frequency) and the temporal modulation frequency. Positive mSNR values (in dB) for a specific recording channel and modulation response subband indicate that speech-synchronized activity has greater power than the background-synchronized activity, whereas negative values indicate that the background-driven synchronization dominates. As with the mMPS, the mSNR patterns vary extensively across background sounds but are often qualitatively similar between recording sites (Fig. 5).

For the eight-speaker babble, mSNRs are near zero across all channels and modulation subbands, indicating that speech-driven power was near equal to the background-driven power. However, for fire and white noise, regions with high mSNR are observed for the modulation subbands in the rhythm range (<16 Hz), consistent with enhanced encoding of slow fluctuations in speech. Yet, for these same recording sites and backgrounds, broad regions of the upper subbands (>16 Hz) had negative mSNRs, suggesting diminished encoding in the voice periodicity range. Note that although the mSNR results across conditions are highly diverse, the acoustic SNR is the same for all conditions shown here (+3 dB). Unlike the acoustic SNR, this multi-channel neural SNR representation suggests that different background sounds may interfere with the neural representation of speech to different degrees, and that this interference may differ depending on the channel and modulation subband.

### Spectrum and Modulation Statistics of Natural Backgrounds Differentially Impact Neural Representations of Speech

Given the substantial differences in the spectrum and modulation statistics of each of the interfering background sounds used here, we next aimed to characterize how these two statistics influence the neural representation of speech in ICC. The differences between the OR, SM, and MM mSNRs for each background indicate that each of these statistics can alter the encoding of speech (Fig. 5). For instance, when the modulation statistics of eight-speaker babble are whitened (SM condition), there is an enhancement in the mSNR for slow fluctuations <16Hz. This suggests that the modulation statistics of speech babble, which have slow fluctuations similar to clean speech (Fig. 3), lead to a distorted speech representation within the low modulation subbands. Conversely for fire, the SM condition is more similar to the OR condition, suggesting that the modulation statistics do not produce as much distortion. Similarly, the differences between the OR and MM conditions show variability in the degree of distortion driven by the background sound spectrum statistics. The bird babble background shows greater distortion in the MM case compared to OR, while the fire and white noise backgrounds show relatively little difference between MM and OR.

By aggregating the results from individual recording sites according to the characteristic frequencies of the sites’ multi-units, we generated mSNRs for the ICC populations (Fig. 6A; see Methods). Here we find that different background sounds appear to distort the neural representation of speech to different degrees, with the spectrum and modulations statistics each exerting separate effects. mSNRs vary extensively across background sounds but are also strongly affected by the sounds’ spectrum and modulation statistics since manipulating these acoustic dimensions (MM or SM, respectively) alters the mSNR. Certain sounds, such as OR eight- and two-speaker babble exhibit low mSNRs over most channels and modulation bands, which suggests that these backgrounds broadly mask speech encoding. Other sounds, such as OR water, jackhammer, freeway and fire, produce negative mSNRs for high-frequency subbands, suggesting that detailed temporal cues in the vocalization pitch range are severely distorted. At the same time, some low-frequency subband information in the rhythm range (<16Hz) may be masked only weakly for these backgrounds, since mSNRs are positive in this range. As in the two recording sites above (Fig. 5), manipulating the modulation and spectrum statistics of each background has background-specific impact at the population level. For eight-speaker babble and bird babble, whitening the modulations (SM) again substantially improved the mSNR. Yet other backgrounds, such as water, construction, and jackhammer noise, show minimal change in mSNRs across OR, SM, and MM conditions.

**Figure 6:**
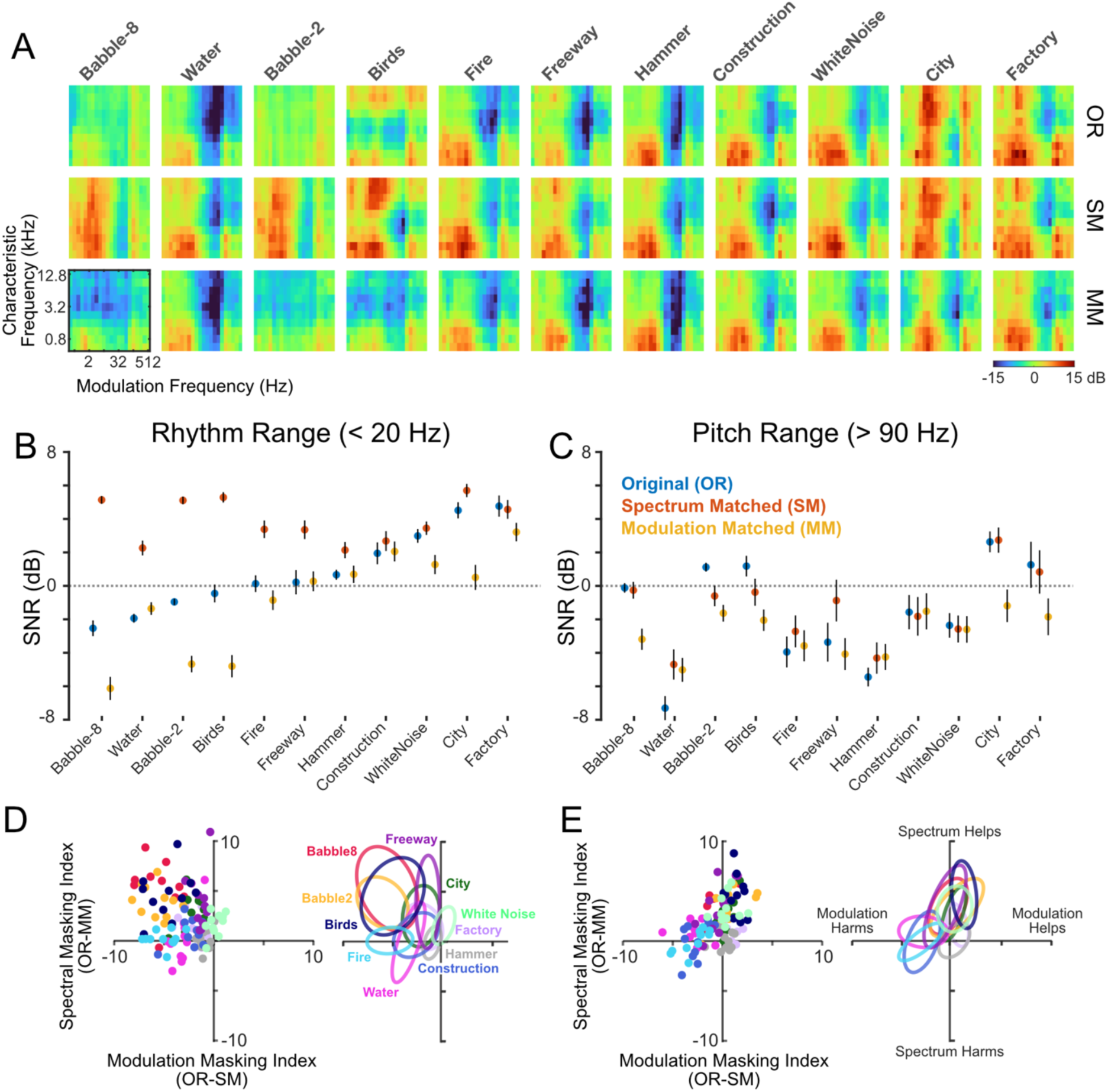
Neural SNR for all background sounds and relative effects of spectrum and modulation statistics. (A) Population-average neural SNR with (n=11) recording sites aligned by characteristic frequency. (B) Average SNR across all channels for temporal modulations in the rhythm range (<20Hz). Dots denote averages across recording sites and error bars denote standard error of the mean, approximated by bootstrapping across sites 1000 times. (C) Average SNR across all channels for temporal modulations in the pitch range (>90Hz). (D-E) Modulation and spectrum masking indices are obtained as the average SNR difference (in dB) between OR and SM vs OR and MM conditions, respectively, and shown for the rhythm (D) and pitch (E) ranges. Dots in (D) and (E) denote individual recording sites. Ellipses in (D) and (E) denote 25% amplitude contours for a bivariate Gaussian distribution model fit for each background.

For each of the backgrounds and perturbations, we summarized the background-specific masking of the speech representation by measuring the population-average mSNR within the rhythm (<20 Hz) and periodicity pitch (>90 Hz) subbands. In general, mSNRs are substantially higher for the rhythm range when compared against the periodicity pitch range of hearing (1.3±3.0 vs -1.9±2.4 dB, mean±SD; average across backgrounds; p<1E-6). This is true even though the overall response power within the periodicity range for clean speech is higher than for the rhythm range (Fig. 2A). Furthermore, although both ranges exhibit background-specific differences resulting from perturbations of the spectrum and modulation statistics, mSNRs for the rhythm range appear to be more variable within a sound category, and thus more strongly impacted by the spectrum and modulation manipulations (MM and SM; within background category variance 9.67 vs. 1.92, p< 0.05). Within the rhythm range, sounds such as the babble sounds (eight-speaker, two-speaker, and bird babble) are strongly impacted by the background modulation statistics, since these sounds tended to have large negative mSNRs for OR and/or SM sounds, when compared to the MM manipulation. Factory, freeway, and white noise, by contrast, were more impacted by the sound spectrum, since manipulating the spectrum (MM) leads to a reduction in mSNR (relative to OR and SM).

To characterize the specific contribution of spectrum- and modulation-based interference, we compute a masking index for each statistic and for each of the rhythm and voice periodicity ranges separately (see Methods). Here, the spectrum masking index (SMI) corresponds to the change in mSNR between OR and MM conditions (matched modulations but different spectra), whereas the modulation masking index (MMI) corresponds to the change in mSNR between OR and SM (matched spectra but different modulation statistics). When calculated for different recording locations, these indices form clusters corresponding to the consistent effects of each background sound on mSNR (Fig. 6 D-E). Within the rhythm range, sounds such as eight- and two-speaker babble clustered broadly at negative MMIs and positive SMIs, indicating that the modulations of these original sounds had a negative impact on the speech encoding. However, within the periodicity pitch range for both sounds, the SMI and MMI both clustered near zero.

For the fire background, MMIs cluster at negative values for both the rhythm and periodicity pitch ranges, suggesting that OR background modulations degrade the representation of speech more so than the SM background within these subbands. Finally, white noise was tightly clustered and had a near-zero MMI (note SM white noise is identical to OR white noise) and a slight positive SMI (for both the rhythm and periodicity ranges), suggesting that the speech encoding deteriorated when the spectrum was manipulated to be 1/f. Collectively, these results demonstrate that different original background sounds produce consistent modulation subband-specific spectrum and modulation masking effects across recording locations, which tend to either improve or degrade the neural representation of speech relative to backgrounds with modified statistics.

Although the mSNR patterns in the ICC show substantial complexity, the neural speech representation may largely reflect acoustic interference at the cochlear level, which in turn reflects interference due to the sound statistics. We thus assess whether the peripheral cochlear model MPS can account for the observed spectrum and modulation masking trends observed in the ICC across background sounds. Here we estimated the cochlear model SNRs (Fig. 7, see Methods), which provide a direct comparison to the mSNRs from the ICC (Fig. 6A). Although differences are visible, the cochlear model SNRs for each of the natural backgrounds and manipulations have patterns similar to the mSNRs. Pointwise scatter plots (across frequencies and modulation subbands) confirm that the cochlear model SNRs are highly correlated with the neural mSNRs across backgrounds and perturbations (OR, SM, and MM), and account for ∼53% of the variance in neural SNRs (OR, *r*^2^ = 0.53 [0.45 0.60]; SM, *r*^2^ = 0.66 [0.57 0.71]; MM, *r*^2^ = 0.58 [0.49 0.66]; mean [95% CI]). This indicates that the ICC neural representation of speech likely reflects, in large part, the frequency- and modulation-specific interference present at the cochlear level.

**Figure 7:**
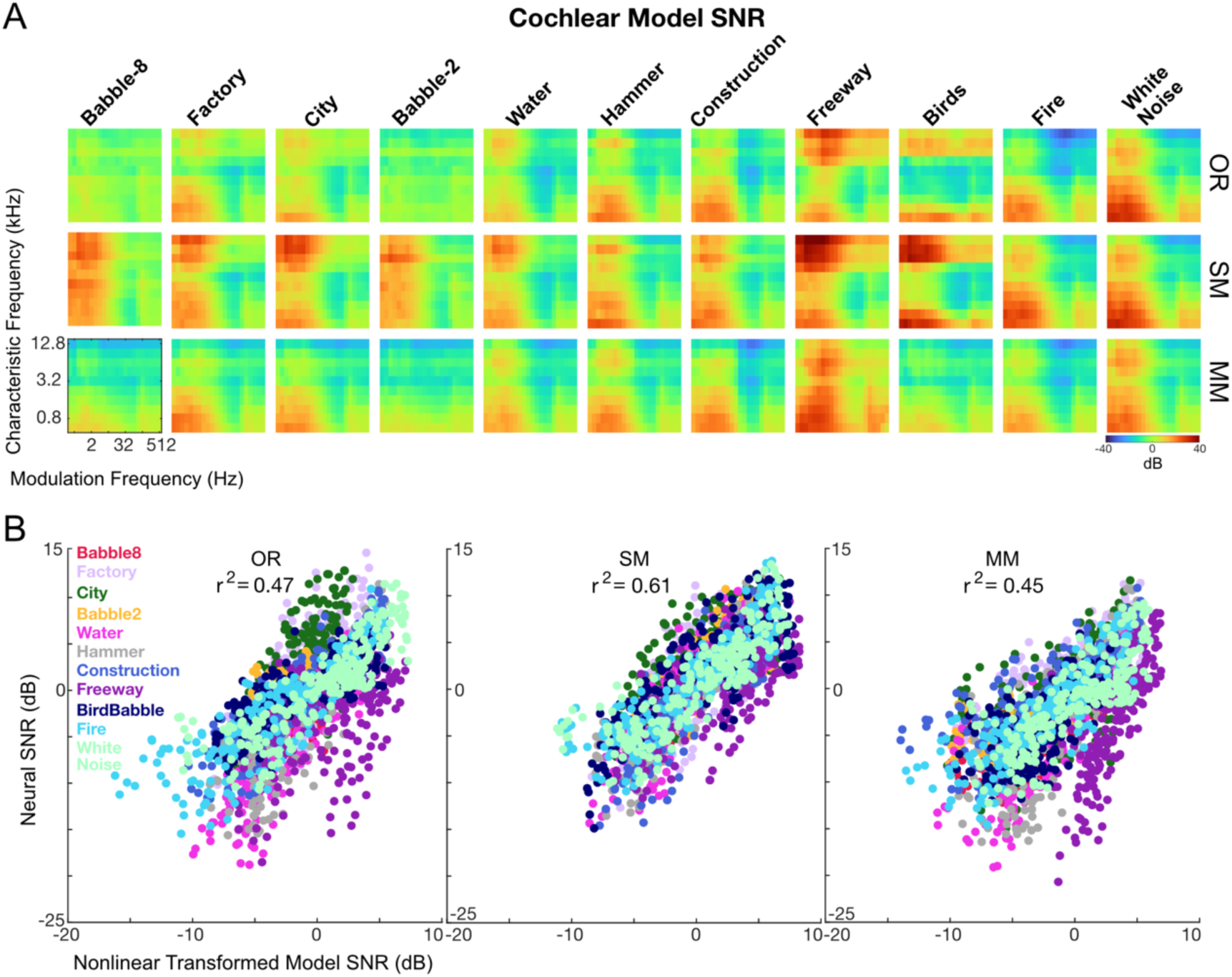
Cochlear model SNR correlates with experimentally-observed mSNR from ICC recordings. (A) Cochlear model SNR shows background-dependent variability in SNR similar to mSNR patterns in ICC (Fig. 6). (B) Additionally, cochlear model SNR is highly-correlated with the neural SNR (for different backgrounds and conditions: OR, SM, and MM).

### Acoustic SNR contributes to speech masking independently of spectro-temporal statistics

Previous perceptual results with human listeners suggest that the acoustic SNR (aSNR) influences perception of speech independently of the modulation statistics of the background sound ^10^. In an additional series of recordings, we thus tested whether the acoustic SNR and the modulation statistics of each background independently impact the neural representation of speech in ICC. Neural mSNRs were measured for speech mixed with OR background sounds across a range of acoustic SNRs (-9 to +3 dB). Population mSNR patterns not only vary considerably across different backgrounds, characteristic frequencies and temporal modulations (Fig. 8A; similar to Fig. 6), but they also vary in a graded fashion across aSNRs. In general, lower acoustic SNRs produce lower neural mSNRs for both the rhythm and periodicity pitch ranges, with the background-specific spectral and modulation masking patterns appearing to be conserved across aSNRs (Fig. 8A).

**Figure 8:**
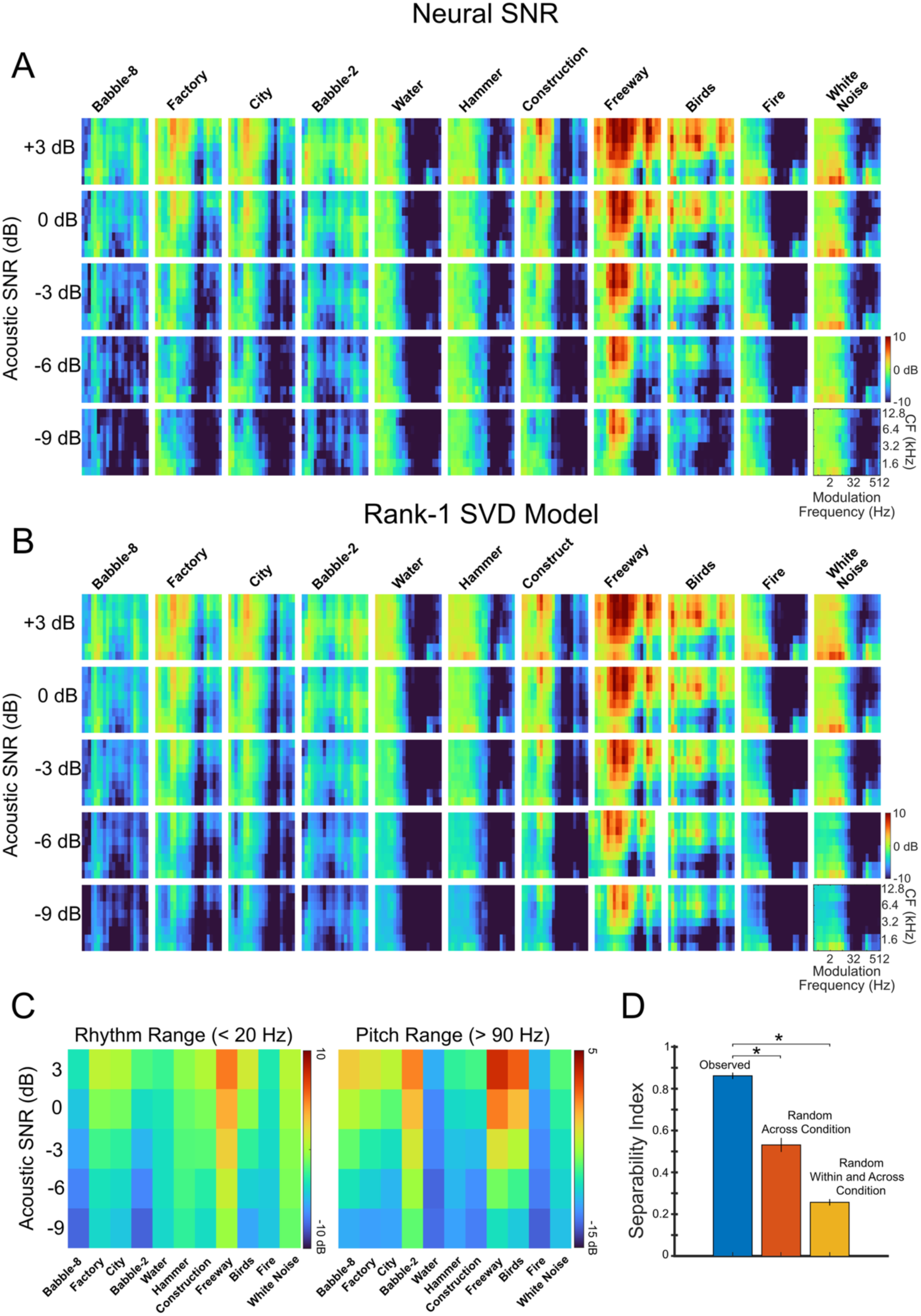
Separability of neural SNR measured at multiple acoustic SNRs. (A) In additional experiments, we measure population-average mSNR in ICC at acoustic SNRs of -9, -6, -3, 0, and 3 dB (speech mixed with OR backgrounds only; n=14 sites). (B) The patterns across acoustic SNRs are well-described by a separable rank-1 model. (C) The rank-1 pattern is visible when averaging within the rhythm (<20 Hz) and pitch ranges (>90 Hz). (D) Separability of the measured mSNR is significantly larger than for shuffled data (bootstrap t-test, * indicates, p < 0.001; red=shuffle backgrounds and sSNR; yellow= shuffle backgrounds, sSNR, frequencies, and modulations).

To determine whether aSNR and background contribute independently to neural masking, we reconstruct the observed mSNRs with a separable rank-1 model in which a single fixed frequency and modulation subband mSNR pattern for each background is simply scaled by an aSNR-dependent gain factor common across backgrounds (see Methods). The goodness of fit for this reconstruction (Fig 8B) then provides a separability index that indicates whether the effect of modifying the aSNR is uniform across background sound categories (see Methods). The measured separability index is high (SI = 0.86±0.01) which suggests that aSNR and background statistics contribute independently to the mSNR patterns. For comparison, we also measure SI in two shuffled datasets, one where the mSNR patterns are randomized across backgrounds and acoustic SNRs, and one where the mSNR patterns are shuffled across backgrounds, acoustic SNRs, frequency channels, and modulation subbands. Shuffling significantly reduces the SI (mSNRs randomized across backgrounds and acoustic SNRs: SI = 0.53±0.03; mSNRs randomized across backgrounds, acoustic SNR, frequency channels, and modulation subbands: SI = 0.26±0.01; bootstrap t-test, p<0.001), suggesting that the observed mSNR patterns scale predictably with acoustic SNR and do not have substantial additional interactions or aSNR-dependent nonlinearities. Rather, similar to human perception ^10^, aSNR and the statistics of each individual background sound appear to contribute independently to masking patterns in ICC.

## Discussion

The results here demonstrate that natural environmental noises interfere with the neural representation of speech in the rabbit ICC in a frequency- and modulation-specific manner. At this level of coding, speech and backgrounds are simultaneously represented, producing a neural interference pattern that reflects the statistics of both sounds. The spectra and modulations present in different natural backgrounds lead to distinct interference patterns, particularly for both slow rhythmic fluctuations at timescales of phonemes and words, as well as faster temporal modulations within the voice periodicity range. Although there is substantial heterogeneity in ICC speech encoding across backgrounds, even when the acoustic signal-to-noise ratio is constant, these patterns are well-described by cochlear model outputs and are conserved even when the acoustic SNR is varied. Altogether, these findings demonstrate that the distortion of speech representations in the rabbit ICC is background- and channel-specific, and that these same patterns may underlie differences in human speech perception during everyday listening.

## The role of Inferior Colliculus

The inferior colliculus is a key auditory hub for processing spectro-temporal sound cues. It is the principal lemniscal midbrain nucleus and the first structure in the ascending auditory pathway to exhibit a preponderance of modulation-selective neurons ^23^. Activity in the ICC is well-described as producing a multi-dimensional decomposition of sounds into frequency and modulation components ^37,38^. Since the modulation statistics of sounds are critical for natural sound recognition and speech perception ^10,29,31,39^, one major function of the ICC may be to set the stage for high-level sound processing by producing an efficient feature decomposition ^34^, which can later be used by downstream cortical regions.

Many previous studies have characterized the neural representation of frequency and modulation cues in the inferior colliculus with well-controlled, synthetic sound stimuli ^18,40–42^. Studies have also identified how the representation of such sounds is altered by some types of noise in IC ^43–45^, such as the representation of tones or modulated signals in white noise. However, there are relatively few studies that have characterized the representation of natural sounds or speech in the IC ^19,24,46^, and none, to our knowledge, have examined speech encoding under such a wide range of natural backgrounds. One unique feature of speech is that it contains detailed spectro-temporal fluctuations or temporal fine structure, and auditory nerve fibers synchronize to such fluctuations up to and exceeding 1 kHz. This synchrony provides a fine-grained representation of speech ^47,48^ that is mostly preserved in the ICC, and which can support neural discrimination of speech sounds ^46,49,50^. Here, we further demonstrate that ICC neural population responses differentially preserve both slow fluctuations at timescales of phonemes and words, as well as faster fluctuations within the voicing periodicity range, under a wide variety of noise conditions.

### Spectrum and Modulation Interference Framework

The findings provide the first neural demonstration of modulation masking in the central auditory system and suggest that the extent to which environmental sounds interfere with the neural coding of speech depends on the frequency channel and modulation subband. Consistent with experimental and theoretical findings^25,34^, ICC exhibits a whitened and presumably more efficient representation of speech when compared to a cochlear model (Fig. 3). Yet, the interference created by environmental noises is well-predicted by the modulation power spectrum of the speech-in-noise mixture as measured through a cochlear model representation. Furthermore, speech and environmental noise appear to both be represented in the neural population activity of ICC. This contrasts with several previous studies that have reported sparse, segregated representations of speech or vocalizations in competing noises in auditory cortex ^22,45,51,52^, where responses are noise- invariant. Other studies using diverse sets of natural sounds, by comparison, reported responses in auditory cortex that are background-dominant ^53^. Although it is plausible that foreground sounds are segregated from competing background noises at some stage, these “noise-invariant” neurons in cortex ^22^ were observed using background sounds with only limited diversity. Additional studies may be needed to determine whether such findings generalize across backgrounds with more varied spectrum and modulation statistics. Here, even at a fixed acoustic SNR, the neural SNRs vary extensively. This suggests that specific speech features can be encoded with low or high fidelity, depending on the recording site’s characteristic frequencies and the specific background sound. A single neuron could thus potentially be masked or unmasked in any given condition, producing responses that are dominated by the foreground or the background or that give the appearance of “noise invariance”. Controlling for the localized SNR directly overlapping the neuron’s receptive field will likely be necessary to confirm how selective the responses in cortex are to general foregrounds or backgrounds.

The results here indicate a complex but consistent ICC representation of speech in noise; however, there are several limitations to these results. First, with only 4 speech and 4 background excerpts lasting 3 seconds each, there could be small, residual contributions of common background features in the estimated foreground-driven response, and likewise of common foreground features in the background-driven response. Excerpts were selected independently and optimized for the typical limited recording times, but, since these are natural sounds, there is some shared structure between excerpts. This residual power is small relative to the driven power, but future recordings using additional excerpts could identify how quickly the residual power decays as the number and duration of excerpts increases. Secondly, an additional limitation may be our focus on population aMUA signals. These signals are substantially less variable than spikes and produce uniform population data covering the tonotopic axis of ICC. However, since these signals average over pools of ICC neurons, they may not fully reflect the computations that occur within spike trains. Future studies may be necessary to fully describe the heterogeneity in mSNR effects at the level of single neurons.

### Relation to Human Speech Perception in Natural Noises

Our results on the neural encoding of speech in noise may also have implications for understanding human perception ^10,27^. A previous study measured speech recognition in human listeners using the same set of backgrounds^10^, and found that the spectra and modulations of the competing natural noises serve as separate cues that can lead to masking or unmasking of speech. Additionally, in human listeners, different modulation subbands appeared to provide different benefits for speech recognition in noise. The psychoacoustic results mirror the broad findings here on neural representations and support the idea that slow rhythmic fluctuations in speech (at timescales of words and phonemes) and fast temporal fluctuations (related to voice periodicity) may be distinct in their neural encoding.

Results in humans also suggest^10^ that human speech recognition behavior may rely on weighting spectrum and modulation cues to account for channel- and background-specific interference. Here we observed that different backgrounds create distinct patterns of interference in the ICC, which can produce distortions of the speech for a select subset of features while producing minimal distortions for other components. The overall variation in neural SNRs across backgrounds (e.g., babble vs white noise) only approximately mirrors the variation in speech recognition accuracy in humans with these same backgrounds. Since the rabbits are passively listening and may not attend to or recognize speech, the neural SNRs measured here likely mirror only the early stages of human speech recognition. An ideal observer could presumably use frequency and modulation information by selectively focusing on subsets of cues with a high neural SNR to enable speech segregation. However, human listeners also use attentive listening and more dynamic cue weighting to improve recognition performance ^54^.

Finally, one potentially important point of alignment between the results here and studies with human listeners is that they both suggest that background statistics and acoustic SNR are independent. In human speech recognition, recognition accuracy varied widely depending on the spectro-temporal statistics of the noise, but changing the acoustic signal-to-noise ratio scaled performance consistently and monotonically across backgrounds^10^. For the neural interference patterns measured here in ICC, we similarly find that although the patterns vary widely with the statistics of the noise, the effect of acoustic SNR is largely independent of these statistics. The consistency in the neural coding of speech in noise in the ICC across aSNRs may thus partially explain the consistency in speech recognition. Altogether, since deficits in speech-in-noise perception are a major concern for individuals with hearing loss and auditory processing disorder, additional work identifying the neural representation of these sounds may ultimately lead to clinical benefits.

## Methods

### Animal Welfare and Institutional Approvals

Experiments were carried out in five Dutch Belted rabbits (4 females and 1 male, ages: 0.5 - 4 years, weights: 1.5 - 3 kg). Animals listened passively to natural sound mixtures while neural population activity was recorded from the auditory midbrain (inferior colliculus). All experiments were approved by the University of Connecticut Animal Care and Use Committee in conjunction with guidelines issued by the National Institutes of Health and the American Veterinary Medical Association.

### Animal Procedures and Surgery

Animals first underwent 1-2 weeks of acclimation during which they were trained to sit still while restrained in a harness wrapped snugly around their torso and limbs. Following the acclimation period, animals underwent the first of two aseptic surgical procedures.

Following sedation with acepromazine, an incision was made along the midline of the scalp between bregma and a point approximately 2-3 mm posterior to lambda. A surgical plane of anesthesia was maintained throughout the procedure using isoflurane (1-3%) and oxygen (1 L/min), and vital signs (electrocardiogram, breathing rate, and temperature) were monitored throughout the surgery. Stainless steel screws and dental acrylic were used to implant a brass bar (for head-fixation) atop the rabbit’s head along the anterior-posterior axis just left of the midline. Additional dental acrylic was used to create a well around exposed skull overlying the right hemisphere between the points of bregma and lambda where neural recordings are performed. Next, cotton balls were inserted into the rabbit’s ear canals to block and protect the eardrums, and vinyl polysiloxane molding material (Reprosil) was subsequently injected into the ear canals. Once the material hardened, it was carefully removed and used to create custom ear molds fit to each rabbit’s ear canals, through which sound mixtures were presented during experimental recordings.

Following a minimum of five days of recovery post-surgery, animals again underwent an acclimation period of up to two weeks during which they learned to sit still while body-restrained in their harness and head-fixed via clamping their head bar. During acclimation, rabbits were gradually introduced to foreground-background mixtures presented via binaural speakers connected to tubes projecting through the ear molds, custom-fit to the rabbit’s ear canals.

Post-acclimation, animals underwent a second surgery during which a ∼4x4 mm craniotomy was performed over the right cerebral hemisphere, with the craniotomy located within the acrylic well and centered about 12-13 mm posterior to bregma. The brain was sterilized and disinfected with antiseptic solution (chlorhexidine), and vinyl polysiloxane impression material (Reprosil) was then poured into the acrylic well and allowed to harden to seal off the sterile craniotomy site.

### Sound Delivery

Sounds are delivered to rabbits via a closed binaural speaker system using DT77 drivers (Beyer Dynamics) connected to tubes and ear molds (custom-fitted for each rabbit) that deliver the sounds directly into the external auditory meatus and adjacent to the eardrum ^55^. Sounds were delivered either via a Tucker Davis Technologies (TDT) RZ6 multifunction processor (tones and noise; 98 kHz sampling rate) or via RME Fireface 800 professional audio card (all natural sounds; 44.1kHz sampling rate). The system was calibrated for each animal, ensuring a flat spectrum within the range of 0.1 to 24 kHz, +/- 3 dB, using a finite impulse response inverse filter implemented in the TDT RZ6 multifunction processor.

During each daily session, we first evaluated whether the recording location was likely to be in the central nucleus of inferior colliculus as defined by a clear dorso-ventral tonotopic gradient ^28^. For each penetration site, we delivered a random sequence of tone-pips (0.1 to 20 kHz; 15 to 85 dB SPL; 50 ms duration with 5 ms cosine ramp; 300 ms inter-tone interval) to estimate frequency response areas for each recording channel. We then used an automatic procedure to determine each channel’s best frequency (BF) and characteristic frequency (CF) ^56^. Next, we delivered the experimental sounds consisting of mixtures of speech and either natural or perturbed background noises. Penetration sites that had a clearly defined dorso-ventral tonotopic gradient, consistent with the central nucleus of IC, were then selected for further analysis.

### Mixtures of Speech and Natural Environmental Noise

Eleven original background sounds were used in our experiments: two-speaker babble (Babble-2), eight-speaker babble (Babble-8), bird babble (Birds), construction noise (Construction), city noise (City), factory noise (Factory), crackling fire (Fire), freeway noise (Freeway), hammering noise (Hammer), running water (Water), and a white noise control (White Noise). We used these same backgrounds in a recent study exploring human perception of speech in background noise ^10^. Collectively, these categories represent a range of sounds that typically accompany speech in real-world listening environments. These categories are also acoustically diverse and vary considerably in their sound spectra and spectro-temporal modulation statistics, both of which critically shape perceptual abilities ^10^.

To better identify the influence of background spectra and modulations on the neural responses in ICC, we present original (OR) backgrounds as well as perturbed backgrounds, generated by altering either the background spectra or the modulations while preserving the other. In a first perturbation, we phase-randomize the OR backgrounds while preserving the spectral amplitudes (using FFT). These spectrum-matched (SM) backgrounds thus have the same spectra as the originals but have whitened modulation statistics. In a second perturbation, we then reweight the spectral amplitudes (using FFT) such that they take on a 1/f-like relationship between power and frequency, while leaving unperturbed the OR background modulations embedded in the phase information. These modulation-matched (MM) backgrounds have the same modulation statistics as the originals but have whitened spectrum statistics.

For each natural background and perturbation (OR, SM, and MM), we then generated mixtures of speech-in-noise similar to those used previously to study human speech perception in noise ^10^. Four 3-second-long excerpts of each background category were mixed with four excerpts of foreground speech extracted from an audiobook reading (adult male speaker) of Shakespeare’s *Hamlet* ^57^. This paring generates 16 (4 backgrounds x 4 foregrounds) unique foreground-background sound mixtures for each experimental condition (OR, SM, and MM) of each background category (Babble-8, Fire, Water, etc.). Unless otherwise noted, foreground-background mixtures were presented at a fixed acoustic SNR (aSNR) of +3 dB.

### Neurophysiology

Extracellular multi-unit activity (MUA) was recorded from the inferior colliculus (ICC) of unanesthetized rabbits that passively listened to foreground-background sound mixtures. MUA was obtained over the course of ∼1-hr. acute sessions using a 64-channel linear electrode array (NeuroNexus) inserted into the ICC prior to each recording session. A total of 41 penetrations were made: 11 featured speech mixed with OR, SM, and MM backgrounds, 11 presented speech mixed with OR and SM backgrounds, 5 included speech mixed with OR and MM backgrounds, and the remaining 14 included speech mixed only with OR backgrounds across multiple acoustic SNRs.

Acclimation and recordings occurred within a double-walled, sound-attenuated booth (Industrial Acoustic Company). Prior to recordings, custom-made 64-channel linear silicon electrode arrays (NeuroNexus, 60-um inter-channel spacing and 12-mm shaft length) were manually inserted into occipital cortex and subsequently driven via a micromanipulator (Burleigh) to depths of approximately 7 - 10 mm, consistent with the anatomical location of the rabbit central nucleus of the inferior colliculus (ICC). A dorso-ventral CF gradient along the recording array was used to further confirm electrode placement within the ICC^28^. ICC responses to foreground-background mixtures were continuously recorded at a sampling rate of 12 kHz with an RP2 preamplifier and an RZ2 real-time processor (TDT). Recordings were subsequently analyzed in MATLAB (MathWorks) via custom code. Voltage signals recorded on each electrode channel were first bandpass-filtered (325 - 3000 Hz). Next, the response envelope was extracted by computing the magnitude of the Hilbert transform and lowpass-filtering (475 Hz cutoff frequency) to extract signal envelopes. Envelopes were then downsampled to 2 kHz to produce an analog multi-unit activity signal (aMUA) ^58^. Finally, aMUA signals were used to generate multi-trial rasters for each recording channel in response to presentations of each foreground-background mixture.

### Midbrain Modulation Power Spectrum (mMPS)

To determine the fidelity of speech encoding in the auditory midbrain, and to relate the neural activity to the statistics of each natural sound being tested, we developed and calculated the midbrain modulation power spectrum (mMPS). This neural response metric is motivated by acoustic measures of the modulation power spectrum statistics of natural sounds ^29,34,59^. Although these prior studies used Fourier analysis or models of the auditory system to characterize modulation statistics of natural sounds, here we use neural response measurements to directly estimate the modulation statistics of sounds as represented in IC neural population activity. Conceptually, mMPS measures the reliable and synchronous neural response power within IC as a function of both the recording channel (or the sound frequency) and the temporal modulation frequency of each natural sound.

For clean speech, the neural mMPS is estimated from the responses to multiple excerpts and response trials for clean speech. The neural activity from each recording location was first represented as a single-trial aMUA neurogram matrix, *X_km_*(*t*, *i*), which represents the speech-driven neural activity as a function of both time (*t*) and the recording channel number (*i*), and where *k* represents the speech excerpt (1-4) and *m* the trial number. These single-trial, single-excerpt neurograms are then decomposed into-subbands that are designed to isolate neural activity that is synchronized to different sound modulation components:

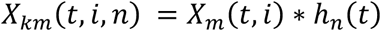

where ℎ*_n_*(*t*) is the impulse response of the n-th subband filter and * is the convolution operator. Subband filters consist of overlapping, logarithmically-spaced, 1-octave-wide b-spline bandpass filters with center frequencies spanning a range of 0.25 - 512 Hz. The subband-filtered neurograms thus capture time-varying neural responses across the recording array at multiple temporal scales (see Fig. 1C). Lower temporal modulation frequency subbands depict coarse patterns of spatio-temporal (or spectro-temporal) activity within IC, while higher modulation frequency subbands depict spatiotemporal response patterns with finer temporal details.

For clean speech excerpt (*k*) and response trial (*m*), the neural response is assumed to consist of a superposition of a stimulus-driven response (*r_k_*(*t*, *i*, *n*)) and an independent neural noise/variability component (*n _m_*(*t*, *i*, *n*)):

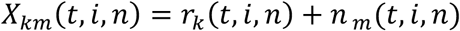

From the single-excerpt and single-trial neurograms for each subband, the clean speech mMPS is derived as the average common response power across trials for each recording channel and modulation subband:

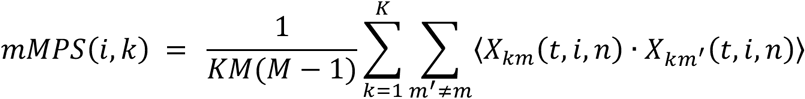

where *K*=4 is the number of speech excerpts used, *M* is the number of response trials (typically 2-4), 〈·〉 is the time-average operator, and where the inner sum represents a pairwise across-trial shuffling of the neurogram responses (sum over *m* and *m*\* such that *m*\* ≠ *m*). Since the neural response noise is independent across trials and also independent of the stimulus-driven response, the trial shuffling cancels out the response noise and guarantees that the time-averaged product of *X_km_* and *X_km_*’ strictly captures stimulus-driven power in the neurogram for each recording channel and subband.

### Estimating Foreground and Background Driven mMPS from sound mixtures

The neural mMPS described above represents the neural response statistics by measuring the power at each recording channel and modulation subband, yet it does so only for sounds presented in isolation. For a sound mixture, it would be ideal if the independent contributions of the foreground and background neural responses could be isolated. Conceptually, two separate mMPSs could be derived for a sound mixture, each accounting for either the foreground- or the background-driven neural activity. Separate foreground- and background-driven mMPS would allow visualization of how neural activity in IC concurrently represents foreground and background sounds and, ultimately, would help determine how background sounds interfere with the neural representation of speech. Below, we propose a trial- and excerpt-shuffling procedure that isolates the foreground (mMPS_F_)- and background (mMPS_B_)-driven mMPS from the mMPS in response to sound mixtures.

We consider a *K* x *L* (*K*=4, *L*=4) set of foreground-background mixtures consisting of speech sentences (the foreground, F; 1 of 4 excerpts per trial) and natural backgrounds (B; 1 of 4 excerpts per trial; e.g., speech babble). The 4x4 structure of the mixtures is leveraged using a shuffling procedure to separate the mMPS into its speech-driven and background noise-driven components. F and B sound excerpts from the mixture are statistically identical across trials and are taken from the same sound recordings, yet they are temporally uncorrelated and independent from each other. For a given F-B mixture, the subband neurogram response is thus a superposition of the foreground- and background-driven components as well as an independent (across trials) additive neural noise:

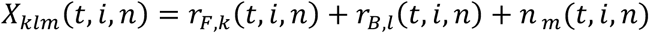

where *r_F_*_,*k*_(*t*, *i*, *n*) is the neural response to the k-th foreground sound, *r_B_*_,*l*_(*t*, *i*, *n*) is the neural response to the l-th background sound, and *n_m_*(*t*, *i*, *n*) is the neural response noise or neural variability on the m-th response trial. By design, the foreground-driven responses, *r_F_*_,*k*_(*t*, *i*, *n*), are independent of the background-driven responses, *r_B_*_,*l*_(*t*, *i*, *n*), and the responses across excerpts are likewise independent (e.g., *r_F_*_,1_(*t*, *i*, *n*) is independent of *r_F_*_,2_(*t*, *i*, *n*)). Thus, in addition to shuffling the neural response across trials to isolate the stimulus-driven component from the neural noise, as in the original mMPS calculation for clean speech, we also shuffle responses across excerpts to isolate the independent contributions of the foreground or background sounds. The foreground-driven mMPS (mMPS_F_) is thus derived by computing the common response power in each recording channel and modulation subband, similar to the original mMPS calculation above:

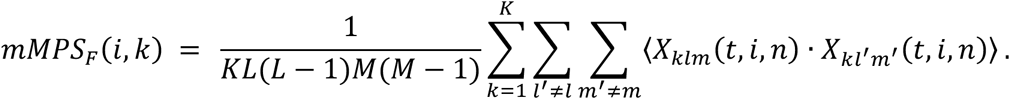

However, in this case, the response power is derived by shuffling across both trials (*m*\* ≠ *m*) and background excerpts (*l*\* ≠ *l*). This mMPS calculation thus isolates the foreground-driven power across each of the recording channels and subbands. Similarly, we can compute the background-driven mMPS (mMPS_B_), but do so by shuffling across foreground excerpts (*k*\* ≠ *k*) and response trials (*m*\* ≠ *m*):

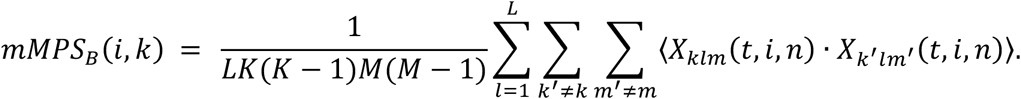

This isolates the background-driven power across each of the recording channels and subbands. Thus, mMPS_F_ and mMPS_B_ account for the response power in the total IC response to a sound mixture that can be attributed to the foreground and background sounds in the mixture.

### Neural Signal-to-Noise Ratio (mSNR)

While the acoustic signal-to-noise ratio is a widely used measure intended to capture the relative interference between a background and foreground sound, the power of natural sounds can vary extensively as a function of both frequency and modulation. Thus, the interference between natural foreground (speech) and background sounds can be both frequency- and modulation-specific, which in turn may impact the neural representation of speech in a frequency- and modulation-specific fashion.

We computed the midbrain neural SNR (mSNR) to assess the unique influences of spectrum and modulation statistics on the fidelity of IC encoding of speech. The mMPS_F_ and mMPS_B_ respectively indicate how strongly IC activity is driven by the speech or background at each recording channel and modulation frequency. The quality of IC encoding of speech can thus be estimated as a neural signal-to-noise ratio (in dB), which is defined as:

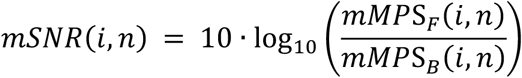

Like the mMPS, mSNR is measured as a function of electrode channel (*i*) and modulation subband (*n*), which captures the impact of frequency (across recording channels) and temporal modulation frequency (across subbands) on the representation of speech. An mSNR > 0 dB indicates that the power of the IC response to speech, at the specific frequency/channel and modulation subband, exceeded that of the response to the background, thus likely enhancing the representation of speech relative to that of the background. Conversely, an mSNR < 0 dB denotes less power in the response to speech compared to the response to the background and implies a weaker representation of speech relative to that of the background.

### Population mMPS and mSNR

To further characterize how the IC neural population represents speech in natural noises, we estimated the population-average mMPS and mSNR. Each recoding site has a unique CF that was derived from the tone-based frequency response area. We thus analyzed the mMPS of clean speech and the mSNR in response to sound mixtures across the population of recording sites (averaged across penetrations and animals) by assigning aMUA responses to ½-octave-wide bins based on the measured CFs (0-8 octaves, relative to 100 Hz; 16 bins between 100 Hz – 25.6 kHz). We then averaged all aMUA responses within each ½-octave CF bin and retained bins with at least 20 multi-units for population-level analysis. The resulting population-average mMPS and mSNR thus represent the speech encoding as a function of frequency (instead of recording channel) and subband/modulation frequency.

### Spectral & Modulation Masking Indices

To further quantify the contributions of the background sound’s spectrum and modulation statistics on the ICC representation of speech, we average across frequency and temporal modulations to compute two distinct masking indices: a spectral and a modulation masking index. These indices are conceptually similar to those used previously to measure spectrum or modulation masking-driven changes in speech recognition accuracy for human listeners. Here, however, we modify these indices and estimate them directly from the neural mSNR measurements. For speech paired with one condition (OR, SM, MM) of one background (fire, water, bird babble, etc.), we first average the mSNR over multi-units within and across channels (i.e., frequency) and a select range of modulation frequencies, thereby yielding one average mSNR for each speech-background mixture for each recording session. The spectral masking index is calculated as the average mSNR difference between speech mixed with an OR background and speech mixed with the MM condition of the same background:

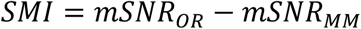

Conceptually, the OR and MM sounds have identical modulation statistics, such that any residual masking that produces a change in neural mSNR must arise from the spectrum statistics. Likewise, the modulation masking index is calculated as the average mSNR difference between speech mixed with an OR background and speech mixed with the SM condition of the same background:

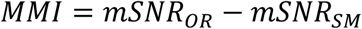

For this scenario, the OR and SM backgrounds have matched spectra, such that changes in mSNR arise because of differences in modulation statistics, which are reflected in the modulation masking index. Since low temporal modulation frequencies are generally associated with the perception of rhythm cues that contribute towards speech recognition, and high temporal modulation frequencies are associated with the perception of voice periodicity and pitch (and are likely more important for voice quality and gender identification), we estimated masking indices separately for these specific ranges: <20 Hz (rhythm) and >90 Hz (voice periodicity). The intermediate range of temporal modulations (∼20-90 Hz), which overlaps with the perceptual range of roughness, was not considered here since it had substantially less power (Fig. 3) and lower average mSNR (-6.9±3.3 dB) compared to the rhythm (1.3±3.0 dB) and pitch (-1.9±2.4 dB) bands.

### Cochlear Model SNR (cSNR)

To characterize the transformation of sound mixtures along the ascending auditory pathway, we use the output of a cochlear model (cochleogram) as a reference for comparing IC neural responses ^34^. The cochlear model takes as input the same speech and background excerpts (speech or background in isolation) outlined above and employs a gammatone filterbank consisting of 0.1-octave-spaced equivalent rectangular bandwidth (ERB) filters (0.02 - 20 kHz). The time-varying sound envelopes in each frequency channel are extracted via the Hilbert transform and then lowpass-filtered at 1000 Hz to simulate the lowpass filtering at the auditory nerve synapse. The resulting cochleogram, *S*(*t*, *f_i_*), represents the envelopes as a function of time (*t*) and the i^th^-frequency channel (*f_i_*).

Next, to characterize the temporal modulations generated at the outputs of the cochlear model and to ultimately estimate the cochlear model SNRs, we performed a subband decomposition of the cochleogram using the same modulation filterbank applied to ICC neural responses:

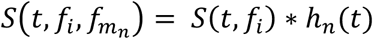

where ℎ*_n_*(*t*) is the filter impulse response of the n-th modulation subband filter (centered around modulation frequency 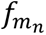; spanning 0.25 - 512 Hz). The cochlear modulation power spectrum of each sound (foreground or background) is then derived by estimating the time-averaged power across each frequency and modulation subband and then averaging across all sound excerpts (*K*=4 backgrounds or foregrounds):

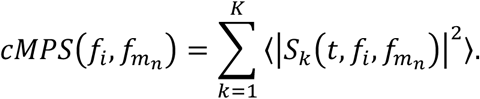

The resulting cochlear model modulation power spectrum (cMPS) depicts the sound power as a function of cochlear channel frequency (*f_i_*) and temporal modulation frequency 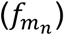, analogous to the neural population mMPS derived above. Finally, we used the foreground and background cMPS to derive the cochlear SNR:

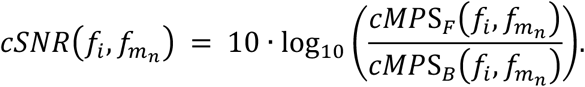

Finally, the cochlear and midbrain SNRs were highly-correlated, despite the cochlear SNR having a substantially larger dynamic range than the neural data, presumably due to neural nonlinearities and noise (such as spiking nonlinearities, neural variability/noise, and compression). We thus applied an affine transformation to rescale the cochlear model SNRs by fitting a second order polynomial to the cSNR vs. mSNR scatter plot and subsequently rescaling the cochlear model SNRs to match the dynamic range of the neural measurements (mSNRs; Figure 7B). Although this resulted in a rescaling of the cSNR dynamic range, this transformation did not alter the relationship between cSNR and mSNR (r=0.75 [0.69 0.80] with vs r=0.75 [0.69 0.80] without, mean [95% CI]).

The cSNR measurements were obtained for speech and each background category (fire, water, bird babble, etc.) and for each of the background conditions (OR, SM, and MM). The cSNR thus provides a measure of acoustic interference (frequency vs modulation) from a cochlear model perspective that can be used to compare directly with the neural mSNR measurements.

### Influences of Acoustic SNR (aSNR) and Modulation Statistics on Masking

To assess the impact of acoustic SNR and modulation statistics of the background sound on the neural encoding of speech, we developed a final acoustic paradigm in which we varied the intensity of our 11 background categories (OR condition only) while keeping the intensity of speech constant. Speech was presented with these variable-intensity OR backgrounds to create mixtures that ranged in acoustic SNR (aSNR) from -9 to 3 dB in 3-dB increments (-9, -6, -3, 0, 3 dB speech-background mixtures as well as clean speech). Using the same experimental setup and procedure as for the previous paradigm, we presented the sound mixtures to rabbits (n=14 recording sites) and estimated mSNR across frequencies and temporal modulations for each speech-background mixture at each aSNR.

To determine whether aSNR and background modulation statistics impact the neural representation of speech independently, we represented the mSNR for each background category and acoustic SNR value as a two-dimensional matrix with rows representing the measured mSNR patterns observed across frequencies, modulation subbands, and background sounds, and columns representing the acoustic SNR (Fig. 8A). We then tested for separability of the effects of aSNR and background modulation statistics on mSNR patterns by applying singular value decomposition (SVD) to generate a rank-1 approximation of the composite mSNR matrix (Fig. 8B). A separability index (SI) was then defined as:

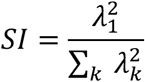

where *λ_k_* is the k-th singular value. Conceptually, the SI represents the fractional power in the mSNR accounted for by the separable rank-1 mSNR approximation, where values close to 1 indicate that the two variables (i.e., aSNR and background modulation statistics) contribute independently to the observed mSNR patterns in IC (patterns across frequencies, modulation subbands, and backgrounds are constant for each sound but are scaled by a aSNR dependent gain). Conversely, values close to 0 suggest that the two dimensions are inseparable and jointly impact the mSNR patterns.

